# CbrA-CbrB Activity is Sensitive to Changes in *Pseudomonas aeruginosa* Cellular Redox State

**DOI:** 10.64898/2026.11.13.766592

**Authors:** Amy Conaway, Jackson C. Shimkonis, Igor Todorovic, Lia A. Michaels, Dhanabala-Subhiksha Rajesh-Khanna, Stacie S. Balsam, Nico Botelho, Jay Goddard, Dallas L. Mould, Colleen E. Harty, Jennifer M. Bomberger, Deborah A. Hogan

## Abstract

In *Pseudomonas aeruginosa*, the CbrA-CbrB two component system contributes to catabolism of diverse carbon and nitrogen sources, several virulence attributes, and is required for the elevated fitness of clinically relevant strains with loss-of-function mutations in *lasR* in passaged cultures. Despite its significance, the mechanism of activation of the CbrA kinase remains unknown. Leveraging the observation that LasR-strains had high CbrA-CbrB signaling that led to increased phosphatase activity, we performed a transposon mutant screen in a LasR-clinical isolate for strains that potentially lost high CbrA-CbrB signaling. Through these studies, we demonstrate that CbrA-CbrB activity correlates with intracellular cysteine abundance and activity is repressed by high or low levels of cellular reducing power. We show that CbrA-CbrB activity is elevated in late-stage cultures and also impacted by stationary phase signaling. Additionally, CbrA-CbrB activity is elevated in the Δ*lasR* mutant in competition with the wild type and this activity is critical for the Δ*lasR* mutant competitive. Feedback inhibition of CbrA-CbrB activity suggests the two-component system plays a role in regulating the cellular redox state. Our findings establish a mechanism by which CbrA-CbrB can activate across diverse conditions and contribute to *lasR* mutant fitness.

**Importance:** In *Pseudomonas spp.,* the CbrA-CbrB two-component system regulates metabolism and several virulence aspects. However, the mechanism of activation of the CbrA kinase remains unknown. We previously found CbrA-CbrB activity is elevated in clinically relevant LasR loss-of-function mutants. Here we leverage this phenotype to perform the first high throughput screen for genes required for elevated CbrA-CbrB activity. This work has revealed that CbrA-CbrB activity is responsive to changes in the cellular redox state and stationary phase signaling. Our findings have laid the groundwork to understand CbrA-CbrB as a global stress response in *P. aeruginosa* and enable the exploration of CbrA as a target for therapeutic intervention.

## Introduction

*Pseudomonas aeruginosa* is an opportunistic bacterial pathogen that can infect multiple body sites in susceptible hosts, including the airways, eyes, chronic wounds, and burns. Loss-of-function (LOF) mutations in the gene encoding the quorum sensing transcriptional regulator LasR are commonly found in both clinical and environmental isolates (1–4). Despite quorum sensing being positively associated with virulence (5, 6), *lasR* mutants are associated with worse clinical outcomes in both acute and chronic infections (1, 7). This paradox has sparked significant interest in understanding the fitness and virulence of *lasR* mutants.

Previously, we found that *lasR* mutants have increased expression of genes associated with virulence and phosphate acquisition at physiological concentrations of inorganic phosphate (Pi) relative to LasR+ comparator strains (8). Virulence and phosphate acquisition are dependent on the two-component phosphate acquisition system PhoR-PhoB. PhoR-PhoB is only active under Pi-limiting conditions and at very low or very high Pi levels, the wild type and Δ*lasR* mutant have similar activity. This indicates that the permissive Pi range for PhoR-PhoB activity is larger in the Δ*lasR* mutants. Importantly, we have found that the elevated PhoR-PhoB activity, increased growth on diverse carbon sources, and dominance in laboratory evolutions associated with *lasR* mutants all require the two-component system CbrA-CbrB (8–10). We propose that the *lasR* mutant cellular environment favors CbrA-CbrB activity and this leads to their increased fitness.

CbrA-CbrB is a global regulatory two-component system specific to and highly conserved within *Pseudomonas spp..* CbrA is a histidine sensor kinase that activates the transcription factor CbrB via phosphorylation. Active CbrB functions with the sigma factor RpoN to induce expression of the non-coding RNA *crcZ*. *crcZ* acts as a sponge, sequestering the translational repression complex Crc-Hfq away from its mRNA targets and allowing for protein expression (see **Fig. 1A** for pathway) (11–14). The Crc regulon is large and highly condition dependent and CbrA-CbrB activity contributes to diverse carbon catabolism (9, 13, 15) and virulence (8, 16, 17). In addition to LasR, the sigma factor RpoS has also been found to repress CbrA-CbrB activity (18). However, the mechanism of CbrA activation remains unknown.

**Fig. 1.**
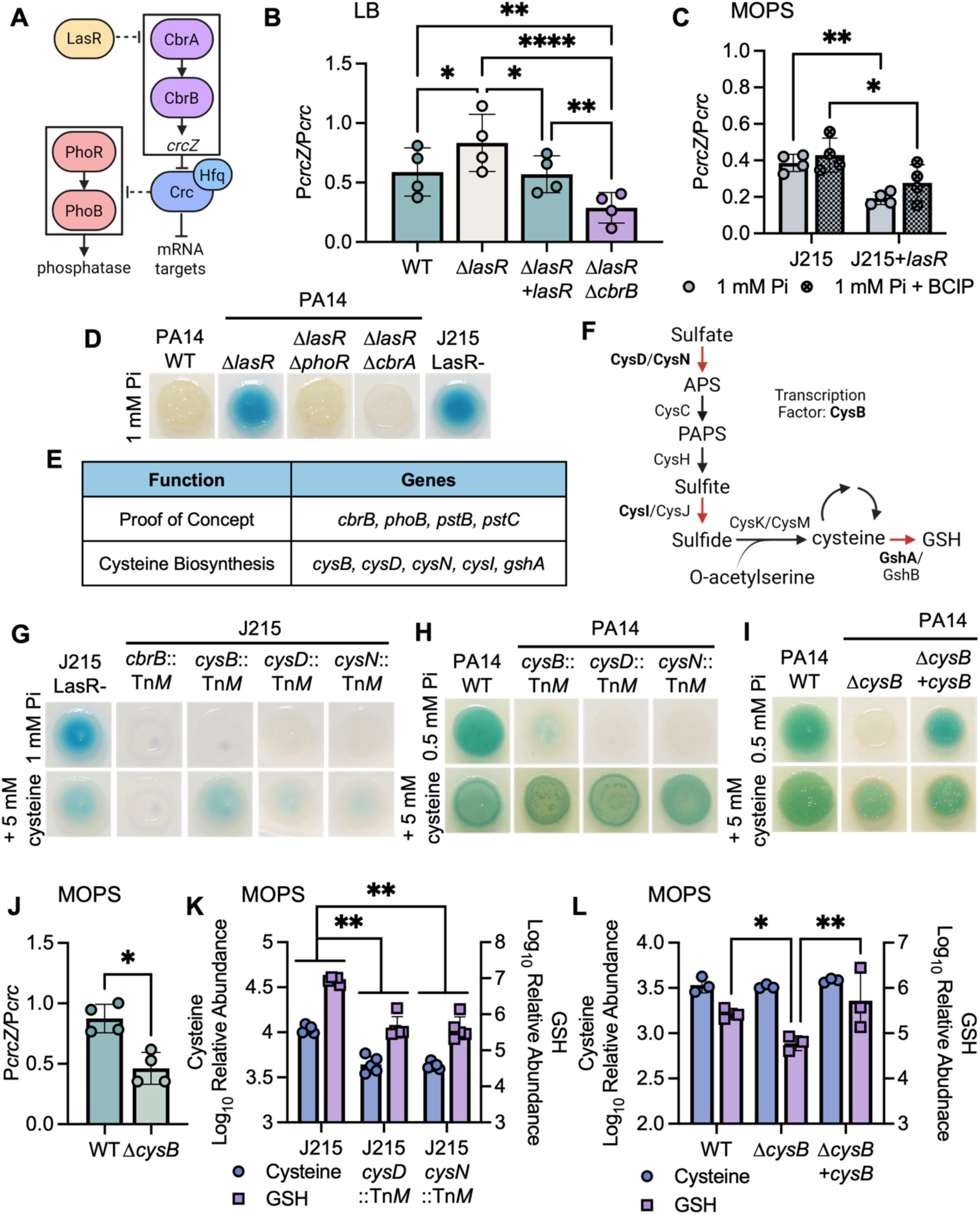
Elevated P*crcZ* activity in *lasR* mutants required cysteine and glutathione (GSH) biosynthesis. **A)** A model of the LasR-CbrA-PhoR pathway. **B)** P*crcZ* activity was significantly elevated in the Δ*lasR* mutant after 16 h growth in LB and repressed to wild-type levels by complementation. Elevated activity required CbrB. Statistical analysis was performed with a repeated measures one-way ANOVA and Tukey’s multiple comparisons to compare each strain (n = 4). **C)** Complementation of the PA14 *lasR* allele in the clinical isolate J215 significantly repressed P*crcZ* activity at 1 mM Pi and the presence of BCIP had no effect. Statistical analysis was performed with a two-way ANOVA and FSD comparing each strain and condition (n = 4). **D)** Representative image of 5 µL spots of *P. aeruginosa* incubated on MOPS-glucose agar with 1 mM Pi and BCIP for 16 h. Blue pigment indicates phosphatase activity. The Δ*lasR* mutant and produced phosphatase and this required *phoR* and *cbrA*. The J215 clincal isolate also produced phosphatase (n = 3). **E)** A random transposon library of the clinical isolate J215 was screened on MOPS-glucose agar with 1 mM Pi and BCIP. Less than 0.5% of mutants screened exhibited altered BCIP cleavage. **F)** Schematic of the cysteine and GSH biosynthesis pathway. Bolded enzymes and red arrows indicate transposon mutants identified in the screen. **G)** Representative images of J215 and mutants grown on MOPS-glucose agar with 1 mM Pi and BCIP, with or without 5 mM cysteine. Blue pigment indicates phosphatase activity. The J215 *cbrB*::Tn*M* mutant was deficient in BCIP cleavage, and this was not rescued by exogenous cysteine. The J215 *cysB*::Tn*M, cysD*::Tn*M,* and *cysN*::Tn*M* mutants were all deficient in BCIP cleavage and rescued by cysteine supplementation (n = 3). **H)** Representative images of PA14 wild type and mutants grown on MOPS-glucose agar with 0.5 mM Pi and BCIP, with or without 5 mM cysteine for 16 h. Blue pigment indicates phosphatase activity. The PA14 *cysB*::Tn*M, cysD*::Tn*M,* and *cysN*::Tn*M* mutants were all deficient in BCIP cleavage and rescued by cysteine supplementation (n = 3). **I)** The Δ*cysB* mutant was deficient in BCIP cleavage on MOPS-glucose agar with 0.5 mM Pi and BCIP and this was rescued by complementation at the native locus or supplementation with exogenous cysteine (n = 3). **J)** P*crcZ* activity was significantly lower in the Δ*cysB* mutant compared to the wild type on MOPS-glucose agar with 0.5 mM Pi. Statistical analysis was performed with an unpaired two-way Welch’s t test (n = 4). **K)** The relative abundance of intracellular cysteine and GSH was significantly lower in the J215 *cysD::*Tn*M* and *cysN*::Tn*M* mutants relative to the parental J215 clinical isolate on MOPS-glucose agar with 1 mM Pi. Statistical analysis was performed with a two-way ANOVA and Sidak’s multiple comparisons test comparing each mutant to J215 (n = 5). **L)** The relative abundance of intracellular GSH was significantly lower in the Δ*cysB* mutant relative to the wild type and complement on MOPS-glucose agar with 0.5 mM Pi. There was no significant difference in cysteine abundance between the strains. Statistical analysis was performed with a two-way ANOVA and Sidak’s multiple comparisons test comparing each strain (n = 3). For all panels * = p < 0.05, ** = p < 0.01, *** = p < 0.001, and **** = p < 0.0001.

Leveraging CbrA-CbrB regulation of phosphatase activity, we developed a high throughput protocol to screen for *lasR* mutants with altered CbrA-CbrB activity. In this paper we show that CbrA-CbrB is repressed by low or high levels of reducing or oxidizing agents. Furthermore, our data suggest that CbrA-CbrB is activated in stationary phase and impacted by multiple stationary phase regulators. Additionally, we found that the Δ*crc* mutant has minimal CbrA-CbrB activity, suggesting that de-repression of Crc targets provides feedback inhibition of CbrA-CbrB signaling. Our findings support a model wherein CbrA-CbrB is activated within a permissive range of redox conditions and Δ*lasR* mutants are more often within this range. We found that forcing Δ*lasR* mutants outside this range by supplying exogenous cysteine represses CbrA-CbrB activity and their competitive fitness. This study has significantly expanded our understanding of CbrA-CbrB regulation and provided a model for activation across diverse conditions. Furthermore, this work provides a deeper insight into the factors contributing to the evolution and increased fitness of *lasR* mutants.

## Results

### The Δ*lasR* mutant had higher levels of CbrB-dependent *crcZ* promoter activity

We previously demonstrated that the *P. aeruginosa* strain PA14 Δ*lasR* mutant has significantly increased expression of the CbrB-regulated noncoding RNA *crcZ* (10) (see **Fig. 1A** for model) (8, 9). To quantify *crcZ* promoter activity, we have adapted an existing transcriptional promoter fusion (18) so that the *crcZ* promoter drives expression of the gene encoding the mOrange2 fluorophore while the promoter of the constitutively expressed *crc* gene fluorophore (11) drives expression of the gene encoding the mScarlet fluorophore; P*crcZ* activity requires CbrB while P*crc* activity does not (**Fig. S1A-B**). For quantification of P*crcZ* activity across strains and conditions, mOrange2 fluorescence (P*crcZ* activity) was normalized to mScarlet fluorescence (P*crc* activity). As expected, normalized P*crcZ* activity was significantly lower in a Δ*cbrB* mutant, and P*crcZ* activity was rescued with complementation with the native *cbrB* gene (**Fig. S1C**).

We observed significantly higher P*crcZ* activity in the Δ*lasR* mutant than the wild type or the complemented strain in LB medium after 16 h and *cbrB* was required for high P*crcZ* activity in the Δ*lasR* mutant. We noted that the Δ*lasRΔcbrB* mutant also had significantly less P*crc* activity than the Δ*lasR* mutant (**Fig. S1D**) in addition to significantly less P*crcZ* activity (**Fig. S1E**), underscoring the importance of normalizing P*crcZ* activity to the reference promoter.

### A genetic screen for mutants with reduced P*crcZ* activity in a LasR-background

We previously reported that at physiological phosphate concentrations, the PhoR-PhoB two-component system, involved in phosphate acquisition, is more active in the Δ*lasR* mutant than in the wild type, and the increased activity requires CbrA-CbrB (see **Fig. 1A** for pathway) (8). We found the elevated PhoR-PhoB activity was consistent in both the PA14 Δ*lasR* mutant and a clinical isolate with a loss-of-function mutation to the gene encoding LasR, J215 (8, 19). We used the dual promoter fusion to quantify P*crcZ* activity in PA14 and J215 strains grown in MOPS-glucose agar with 1 mM Pi with or without BCIP. The J215 isolate had significantly higher P*crcZ* activity than in J215 complemented with a functional strain PA14 *lasR* allele (**Fig. 1C**). The PA14 Δ*lasR* mutant also had significantly higher P*crcZ* activity when compared to the wild type or the *lasR* complemented strain (**Fig. S1F**). P*crcZ* activity was not affected by BCIP. Thus, the elevated CbrB-dependent PhoR-PhoB activity in *lasR* mutants is consistent with elevated P*crcZ* activity.

As we have published, PhoR-PhoB activity can be assessed by monitoring phosphatase-mediated cleavage of the colorless substrate 5-bromo-4-chloro-3-indolyl-phosphate (BCIP) to form an insoluble blue pigment (8, 20). On a medium containing 1 mM inorganic phosphate (Pi), the Δ*lasR* mutant had high phosphatase activity while the wild type did not (**Fig. 1D**). BCIP cleavage required both PhoR and CbrA in the Δ*lasR* mutant. The LasR-clinical isolate J215 also converted BCIP at 1 mM Pi.

Increasing the Pi to 10 mM, a concentration shown to repress PhoR-PhoB activity (8, 21), eliminated phosphatase activity in the Δ*lasR* mutant, but did not affect P*crcZ* activity (**Fig. S1G**). Thus, while Pi concentrations were important for creating a visual readout for CbrA-CbrB activity, low Pi was not required for high P*crcZ* activity in LasR-strains.

We leveraged the high phosphatase activity in *lasR* loss-of-function mutants grown on 1 mM Pi to identify mutants in which CbrA-CbrB activity was altered. We created a library of ∼7,500 Mariner transposon mutants in the J215 strain and screened on MOPS-glucose agar with 1 mM Pi and BCIP. We found that ∼0.5% of mutants had visible differences in BCIP cleavage. Candidate hits were rescreened by testing in triplicate on the same medium using OD_600_-normalized inocula. For those strains that passed the secondary screen, locations of the transposon insertions were identified via arbitrary PCR and Sanger sequencing (see **Supplementary Table 1** for full list of hits). Several of the mutants had mutations likely to affect PhoR-PhoB signaling including an insertion upstream of *phoB* and in the coding sequences of the *pstB* and *pstC* genes, which encode a Pi transporter known to modulate PhoR activity (21–23) (**Fig. 1E**). We also found a Tn*M* insertion in *cbrB* itself, which abolished phosphatase activity at this Pi concentration. These proof-of-concept mutants validate that the screen provided readouts on CbrA-CbrB and PhoR-PhoB activity.

### The transposon mutant screen revealed that cysteine and GSH biosynthesis genes were necessary for elevated CbrA-CbrB activity in a *lasR* mutant

Several transposon mutants with low phosphatase activity had disruptions in genes associated with cysteine and glutathione (GSH) synthesis (**Fig. 1E**) including genes encoding the enzymes CysD, CysN, CysI, and GshA (see **Fig. 1F** for pathway). The screen also identified CysB, the transcription factor that regulates *cysD, cysN,* and *cysI* expression, as necessary for phosphatase activity. Cysteine and GSH are both reducing agents important for balancing the cell’s redox state. We narrowed the scope of our further analysis to the upstream *cys* mutants and excluded the J215 *cysI*::Tn*M* mutant from further analysis due to a clear growth defect. We previously reported that CbrA-CbrB is not necessary for phosphatase activity in very low Pi conditions and the Δ*cbrB* mutant cleaves BCIP at 0.1 mM Pi (**Fig. S1H**) (8). Neither the J215 *cys* mutants nor the J215 *cbrB*::Tn*M* mutant had BCIP cleavage defects when grown on MOPS-glucose agar with 0.1 mM Pi (**Fig. S1I**), indicating a functional PhoR-PhoB pathway.

The findings from our screen suggested that cysteine and GSH biosynthesis modulates CbrA-CbrB activity. We hypothesized that supplementing the cysteine biosynthesis mutants with exogenous cysteine could rescue phosphatase activity. MOPS-glucose agar with 5 mM cysteine was sufficient to restore some phosphatase activity in the J215 *cys* mutants (**Fig. 1G**). This rescue is most successful in the transcription factor mutant *cysB*::Tn*M* while the *cysD*::Tn*M* and *cysN*::Tn*M* mutants were only partially rescued by exogenous cysteine. Exogenous cysteine did not stimulate phosphatase activity in the J215 *cbrB*::Tn*M* mutant suggesting that cysteine affects the pathway upstream of CbrB.

To determine if cysteine biosynthesis affects CbrA-CbrB activity in a LasR+ strain as well, we tested mutants from the PA14 nonredundant transposon collection (24) on MOPS-glucose agar with 0.5 mM Pi, where the PA14 wild type produces phosphatase. Phosphatase activity was absent in Δ*cbrA* and Δ*cbrB* mutants and restored by deletion of the downstream repressor Crc (**Fig. S1J**). Transposon mutants in *cysB, cysD,* and *cysN* had less phosphatase activity in the PA14 laboratory strain and all were rescued by supplementation with 5 mM cysteine (**Fig. 1H**). These mutants had no BCIP defects at 0.1 mM Pi, a concentration at which PhoB was required for phosphatase activity but CbrB was not (**Fig. S1H**). We created a PA14 Δ*cysB* clean deletion mutant and found that like the PA14 *cysB*::Tn*M* mutant, it was deficient in phosphatase activity at 0.5 mM Pi, and this was rescued by complementation at the native locus or supplementation with 5 mM cysteine (**Fig. 1I**). Furthermore, we found that the Δ*cysB* mutant had significantly lower P*crcZ* activity on MOPS-glucose agar with 0.5 mM Pi, consistent with our model that phosphatase activity under these conditions is reflective of CbrA-CbrB activity (**Fig. 1J**).

To further explore the relationship between cysteine and CbrA-CbrB activity, we quantified intracellular cysteine and GSH in colonies grown on MOPS-glucose agar with 1 mM (J215 background) or 0.5 mM (PA14 background) Pi for 12 h. We found significantly less cysteine and GSH in the J215 *cysD*::Tn*M* and *cysN*::Tn*M* mutants relative to their parental strain (**Fig. 1K**). While intracellular cysteine was not significantly different in the PA14 Δ*cysB* mutant compared to the wild type or complement, the Δ*cysB* mutant had significantly less GSH which was restored with complementation (**Fig. 1L**). Thus, disruption of the cysteine biosynthesis pathway resulted in lower CbrA-CbrB activity perhaps due to decreases in the intracellular cysteine and/or GSH pools.

### Cysteine accumulation preceded elevated P*crcZ* activity in *lasR* mutants

Our data were consistent with a model wherein LasR-mutants have elevated cysteine/GSH pools and this promotes CbrA-CbrB activity (see **Fig. 2A** for model). Further supporting this model, we found that after 12 h growth on MOPS-glucose agar the LasR-J215 clinical isolate had significantly more intracellular cysteine and GSH than J215 with a functional PA14 *lasR* allele (J215+*lasR*) (**Fig. 2B**).

**Fig. 2.**
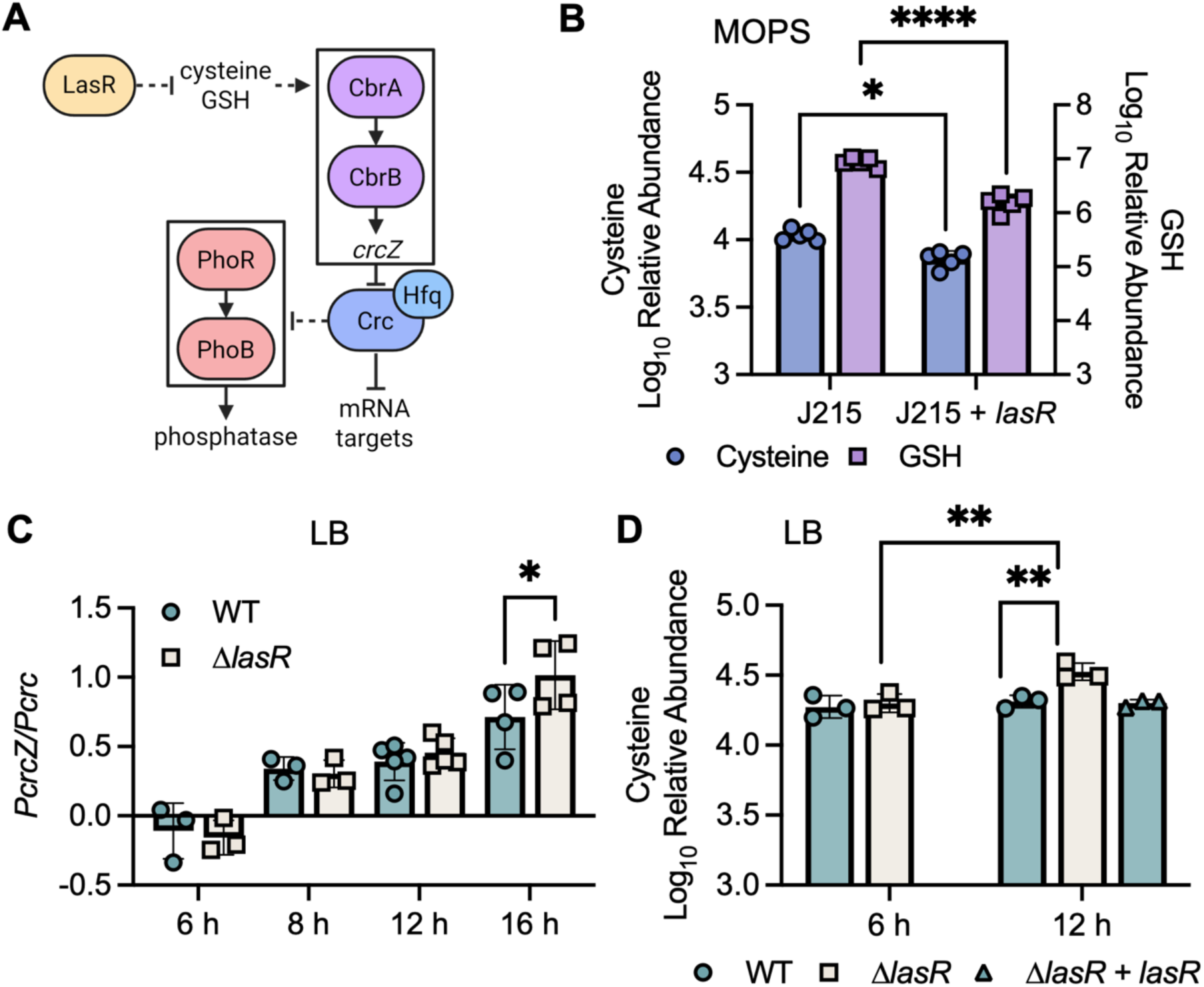
Elevated cysteine levels preceded elevated P*crcZ* activity in the Δ*lasR* mutant. **A)** A model of LasR regulation of CbrA-CbrB activity through changes in intracellular cysteine or GSH pools. **B)** The relative abundance of intracellular cysteine and GSH was significantly lower in J215+*lasR* than the J215 parental strain on MOPS-glucose agar with 1 mM Pi. Statistical analysis was performed with a two-way ANOVA and Sidak’s multiple comparisons test comparing each strain (n = 5). **C)** P*crcZ* activity was below the autofluorescence threshold of the medium (limit of detection) after 6 h of growth in LB medium. Activity was detectable and the same between the wild type and Δ*lasR* mutant at 8 and 12 h (p > 0.54). At 16 h, the P*crcZ* activity was significantly higher in the Δ*lasR* mutant. Statistical analysis was performed with a two-way ANOVA and uncorrected FSD comparing the wild type and Δ*lasR* mutant at each time point (n = 3-5). **D)** The relative abundance of cysteine is the same in the wild type and Δ*lasR* mutant after 6 h growth in LB. Cysteine does not change in the wild type between 6 and 12 h. Cysteine levels are higher in the Δ*lasR* mutant at 12 h compared to the wild type or complement or the Δ*lasR* mutant at 6 h. Statistical analysis was performed with a two-way ANOVA and uncorrected FSD comparing each time point and the wild type and Δ*lasR.* The Δ*lasR+lasR* complement was only tested at 12 h and so excluded from the statistical analysis (n = 3). For all panels * = p < 0.05, ** = p < 0.01, *** = p < 0.001, and **** = p < 0.0001.

To test our model that changes in cysteine/GSH precede CbrA-CbrB activity, we investigated P*crcZ* activity, cysteine, and GSH levels over time in liquid LB cultures. P*crcZ* activity was below the limit of detection due to autofluorescence of LB medium until after 6 h of growth (**Fig. 2C**). Over the course of growth, the wild type and Δ*lasR* mutant were similar until after 12 h, when cultures were in late stationary phase (**Fig. S2A**), when activity increased and was significantly higher in the Δ*lasR* mutant. There was no difference in intracellular cysteine between the wild type and Δ*lasR* mutant at 6 h (**Fig. 2D**), when P*crcZ* activity is undetectable (**Fig. 2C**). Cysteine significantly increased in the Δ*lasR* mutant from 6 to 12 h and was elevated compared to the wild type and the Δ*lasR*+*lasR* complemented strain. GSH did not change from 6 to 12 h and was not significantly different between the wild type and Δ*lasR* mutant (**Fig. S2B**).

We previously found that on LB agar *lasR* mutants have less cysteine and GSH that the wild type (9, 25). Consistent with a connection between cysteine and CbrA-CbrB activity, P*crcZ* activity was significantly lower in the Δ*lasR* mutant relative to the wild type under this condition (**Fig. S2C**). Taken together, our data demonstrate that in liquid LB cultures cysteine levels increased in the Δ*lasR* mutant prior to an increase in P*crcZ* activity and elevated P*crcZ* activity was lost in a condition where the Δ*lasR* mutants have low intracellular cysteine.

### High levels of exogenous cysteine repressed P*crcZ* activity in the Δ*lasR* mutant

To our surprise, supplementing MOPS-glucose agar (0.5 mM Pi) with 5 mM cysteine significantly repressed P*crcZ* activity in the PA14 Δ*lasR* mutant such that it was not significantly different from the wild type or complement (**Fig. 3A**). Exogenous cysteine had the same effect in liquid LB cultures, repressing P*crcZ* activity to wild-type levels in the Δ*lasR* mutant (**Fig. 3B**) without effecting growth (**Fig. S3A**). To gain more insight into the changes in the population with added cysteine, we used confocal microscopy to quantify the P*crcZ* activity in single cells grown in LB liquid medium for 16 h. We found a normal distribution of activity among the cells and that the average activity was elevated in the Δ*lasR* mutant compared to the wild type (**Fig. 3C**). Supplementing the medium with 5 mM cysteine did not affect population-level P*crcZ* activity in the wild type (**Fig. 3B**) but did increase the percentage of cells with higher P*crcZ* activity (**Fig. 3C**). The per cell P*crcZ* activity of the wild type with exogenous cysteine was similar to that of the Δ*lasR*, which had elevated intracellular cysteine (**Fig. 2D**). Exogenous cysteine repressed P*crcZ* activity in the Δ*lasR* mutant both on a population level (**Fig. 3B**) and in single cells (**Fig. 3C**).

**Fig. 3.**
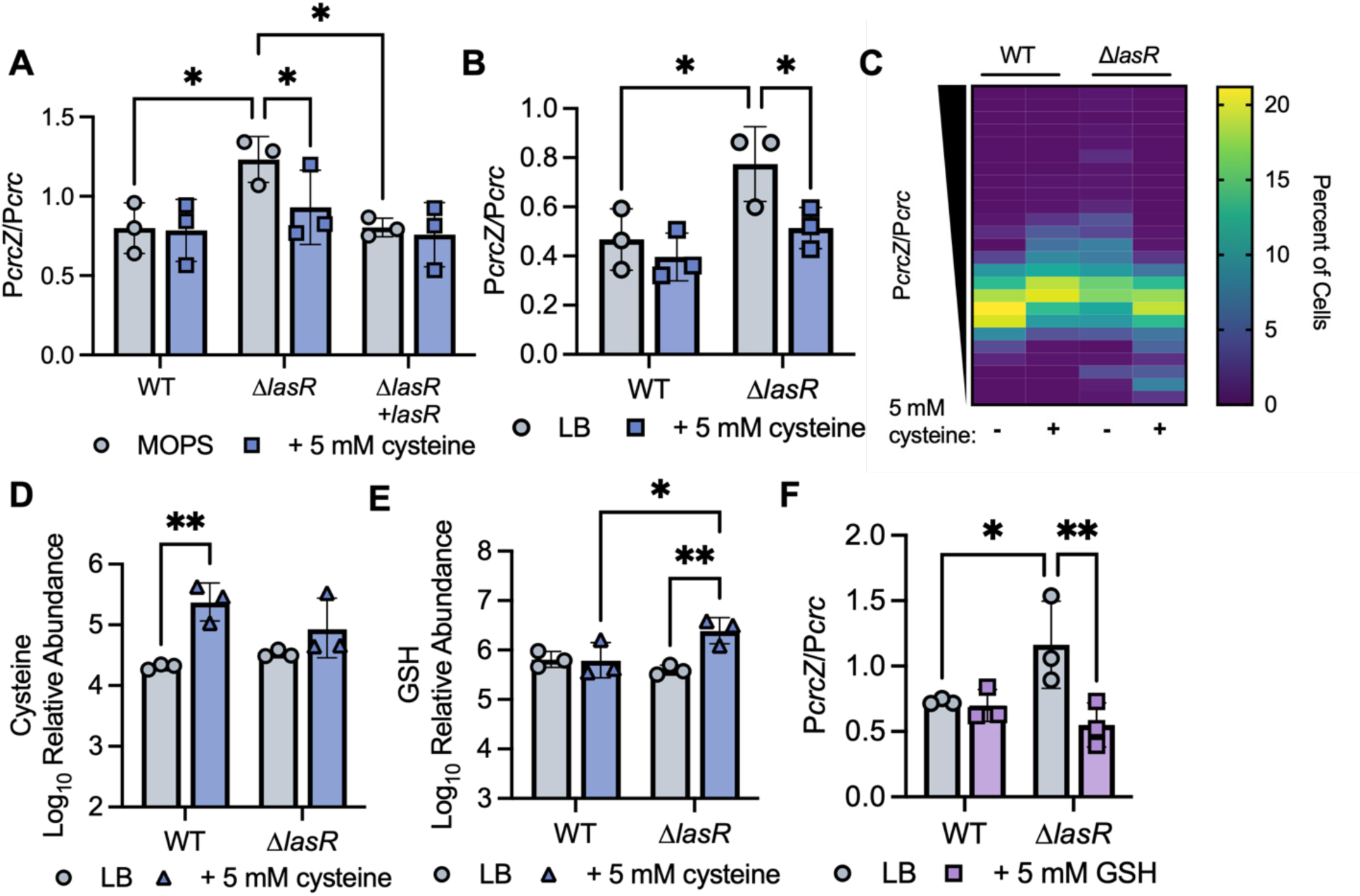
Exogenous cysteine and GSH repressed P*crcZ* activity in the Δ*lasR* mutant. **A)** P*crcZ* activity was significantly elevated in the Δ*lasR* mutant compared to the wild type and Δ*lasR+lasR* complement grown on MOPS-glucose agar with 1 mM Pi. Supplementation of 5 mM cysteine repressed activity such that it was no different from the wild type and complement. Statistical analysis was performed with a two-way ANOVA and Tukey’s multiple comparisons tests to compare each strain and condition (n = 3). **B)** P*crcZ* activity was significantly elevated in the Δ*lasR* mutant grown in LB medium for 16 h. Supplementation with 5 mM cysteine repressed activity down to wild type levels. Statistical analysis was performed with a two-way ANOVA and uncorrected FSD to compare each strain and condition (n = 3). **C)** A heatmap of P*crcZ* activity in single cells after growth in LB for 16 h. Quantification was performed using confocal microscopy of cells on agarose pads. The average P*crcZ* activity was higher in the Δ*lasR* mutant compared to the wild type. The addition of 5 mM cysteine promoted P*crcZ* activity in the wild type and repressed P*crcZ* activity in the Δ*lasR* mutant (n = 3). **D)** Supplementation with cysteine significantly increased intracellular cysteine in the wild type but not the Δ*lasR* mutant (p = 0.12). Statistical analysis was performed with a two-way ANOVA and FSD comparing each strain and condition (n = 3). The LB data shown here is also shown in Fig. 2E. **E)** Supplementation with cysteine significantly increased intracellular GSH the Δ*lasR* mutant but not the wild type (p = 0.93), such that GSH was significantly higher in the Δ*lasR* mutant compared to the wild type. Statistical analysis was performed with a two-way ANOVA and FSD comparing each strain and condition (n = 3). The LB data shown here is also shown in **Fig. S2C**. **F)** GSH significantly repressed P*crcZ* activity in the Δ*lasR* mutant to wild-type levels in LB medium. Statistical analysis was performed with a two-way ANOVA and FSD comparing each strain and condition (n = 3). For all panels * = p < 0.05, ** = p < 0.01, *** = p < 0.001, and **** = p < 0.0001.

To understand the effects of exogenous cysteine on intracellular levels, we quantified intracellular cysteine and GSH with and without supplementing the medium with 5 mM cysteine. The intracellular cysteine levels in the wild type significantly increased with exogenous cysteine supplementation (**Fig. 3D**). However, there was no significant difference in the Δ*lasR* mutant. Instead, the Δ*lasR* mutant had significantly more GSH, while the wild type did not (**Fig. 3E**). This suggested that GSH may repress P*crcZ* activity, as cysteine did, in the Δ*lasR* mutant. This was further supported by the J215 *gshA*::Tn*M* mutant we recovered in the initial screen that lacked phosphatase activity (**Fig. 1E**). We tested the effects of 5 mM exogenous GSH on P*crcZ* activity and found that it repressed activity in the Δ*lasR* mutant to wild-type levels (**Fig. 3F**) without affecting growth (**Fig. S3B**). Taken together, our data demonstrate that while cysteine and GSH biosynthesis were necessary for P*crcZ* activity, high levels of these reducing agents can be inhibitory.

### Carbon limitation promotes P*crcZ* activity in late-stage cultures

Disruption of cysteine biosynthesis inhibited CbrB-dependent phosphatase activity in both the LasR-J215 and the LasR+ PA14 strains (**Fig. 1H-I**). To further explore the effects of exogenous cysteine on P*crcZ* activity in LasR+ and LasR-strains, we grew the PA14 wild type and Δ*lasR* mutant in a cysteine-free minimal medium (M63) with 20 mM glucose. After 16 h growth the Δ*lasR* mutant had significantly more P*crcZ* activity than the wild type or complement (**Fig. 4A**). However, decreasing the glucose to 1 mM elevated the P*crcZ* activity in the wild type and complement such that it was the same as the Δ*lasR* mutant (**Fig. 4B**). In this high P*crcZ* activity condition, both the wild type and the Δ*lasR* mutant were sensitive to P*crcZ* repression by 5 mM exogenous cysteine (**Fig. 4C**) without affecting growth (**Fig. S3C**).

**Fig. 4.**
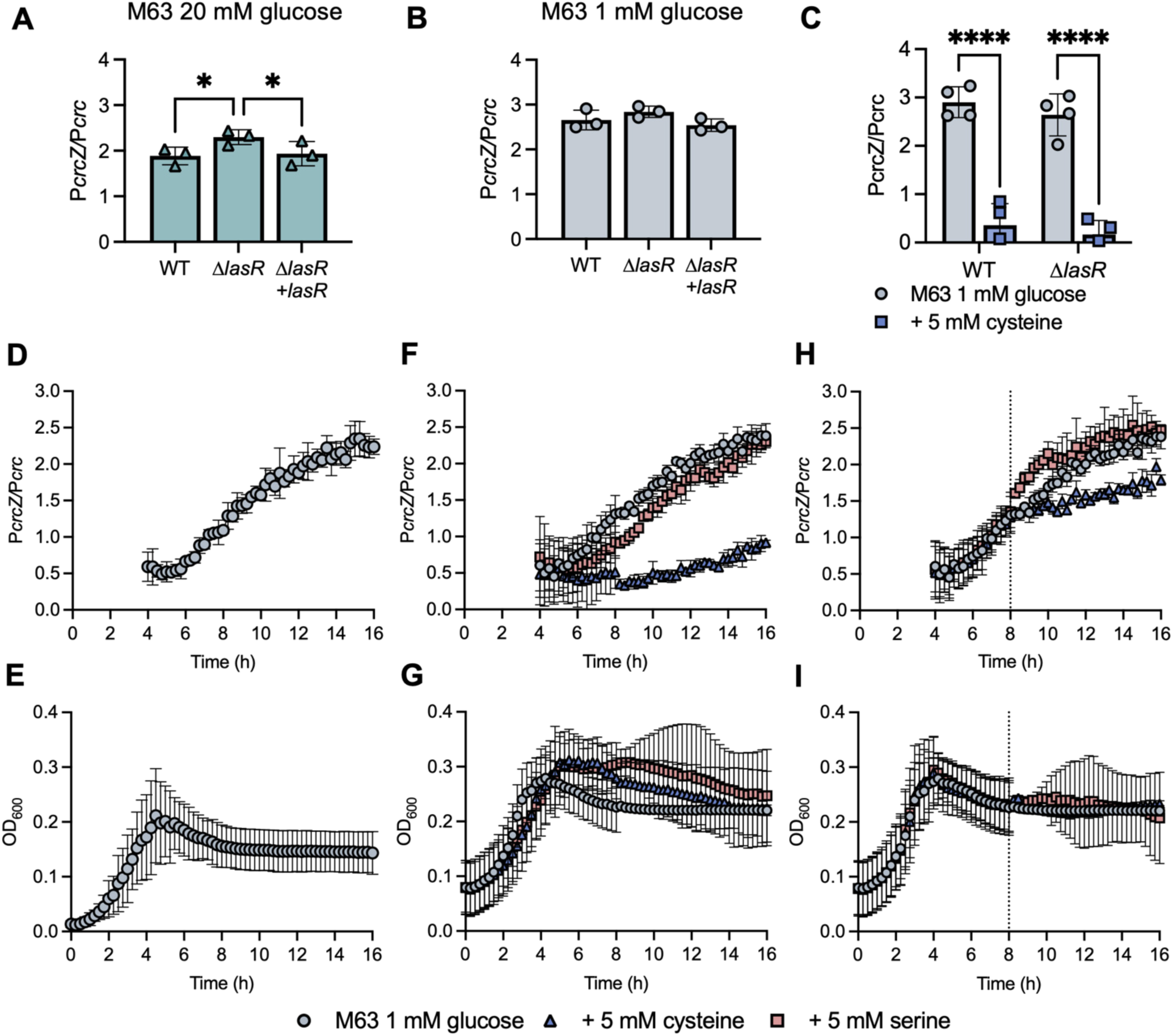
Carbon limitation promoted P*crcZ* activity and cysteine repressed P*crcZ* activity in late-stage cultures. **A-B)** P*crcZ* activity was significantly higher in the Δ*lasR* mutant relative to the wild type and Δ*lasR+lasR* complement grown for 16 h in M63 minimal medium with 20 mM glucose (**A**). There were no differences in activity between the strains when grown in M63 with 1 mM glucose (p > 0.12) (**B**). Statistical analysis was performed with a two-way ANOV and Tukey’s multiple comparisons test comparing each strain in the same condition (n = 3). **C)** P*crcZ* activity was repressed in both the wild type and Δ*lasR* mutant by the addition of 5 mM cysteine when grown for 16 h in M63 medium with 1 mM glucose. There was no difference in activity between the wild type and Δ*lasR* in either condition (p > 0.35). Statistical analysis was performed with a two-way ANOVA and FSD comparing each strain and condition (n = 4). **D)** P*crcZ* activity increased over time in the wild type grown in M63 medium with 1 mM glucose. Data points before 4 h are not shown due to high variability as the fluorescent signal from the overnight culture used for inoculation decreased down to baseline (n = 3). **E)** The wild type grew for ∼ 5 h before reaching maximum density and entering stationary phase in M63 medium with 1 mM glucose (n = 3). **F)** P*crcZ* activity in the wild type grown in M63 minimal medium with 1 mM glucose was repressed by the addition of 5 mM cysteine and unaffected by 5 mM serine (n = 3). **G)** Supplementing 5 mM cysteine or serine to M63 minimal medium did not alter growth (n = 3). **H)** P*crcZ* activity in the wild type grown in M63 minimal medium with 1 mM glucose was repressed by the addition of 5 mM cysteine and unaffected by 5 mM serine after 8 h growth (dotted line indicates when amino acids were supplemented) (n = 3). **I)** Supplementing 5 mM cysteine or serine to M63 minimal medium after 8 h growth did not alter growth (dotted line indicates when amino acids were supplemented) (n = 3). For all panels * = p < 0.05, ** = p < 0.01, *** = p < 0.001, and **** = p < 0.0001.

### Effects of exogenous cysteine on P*crcZ* activity in late-stage cultures

We leveraged the elevated *PcrcZ* activity in the wild type grown in M63 with 1 mM glucose to investigate the effects of exogenous cysteine on early and late-stage cultures. Cultures grown in 1 mM glucose activated P*crcZ* after entering stationary phase at ∼ 5 h, which was amenable to time-course assays (**Fig. 4D-E**). The presence of 5 mM cysteine in the medium at the time of inoculation delayed and attenuated the induction of P*crcZ* activity (**Fig. 4F**). We used serine as a control as both amino acids can be used as carbon sources but only cysteine repressed P*crcZ* activity. The addition of either amino acid had no effect on growth (**Fig. 4G**). Supplementing the medium with 5 mM cysteine after 8 h of growth, when cells were in stationary phase, repressed further induction of P*crcZ* activity (**Fig. 4H-I**). The addition of serine at 8 h had no effect. We tested other metabolites in the cysteine biosynthesis pathway (SO_4_, serine, and methionine) and found that only cysteine significantly changed P*crcZ* activity and none affected growth (**Fig. S4A-B**).

### The effects of reducing and oxidizing agents on P*crcZ* activity

We demonstrated that both low (**Fig. 1H-I**) and high (**Fig. 4C**) levels of cysteine repressed P*crcZ* activity in LasR+ and LasR-strains. We proposed the intermediated amounts of cysteine could promote P*crcZ* activity and so incubated the PA14 wild type in M63 with 1 mM glucose and 0 – 10 mM cysteine (**Fig. 5A**). Intermediate amounts of cysteine (0.625 mM) elevated P*crcZ* activity while 10 mM significantly repressed activity (**Fig. 5C**). The cells grew well in all conditions, indicating these levels of cysteine are not toxic.

**Fig. 5.**
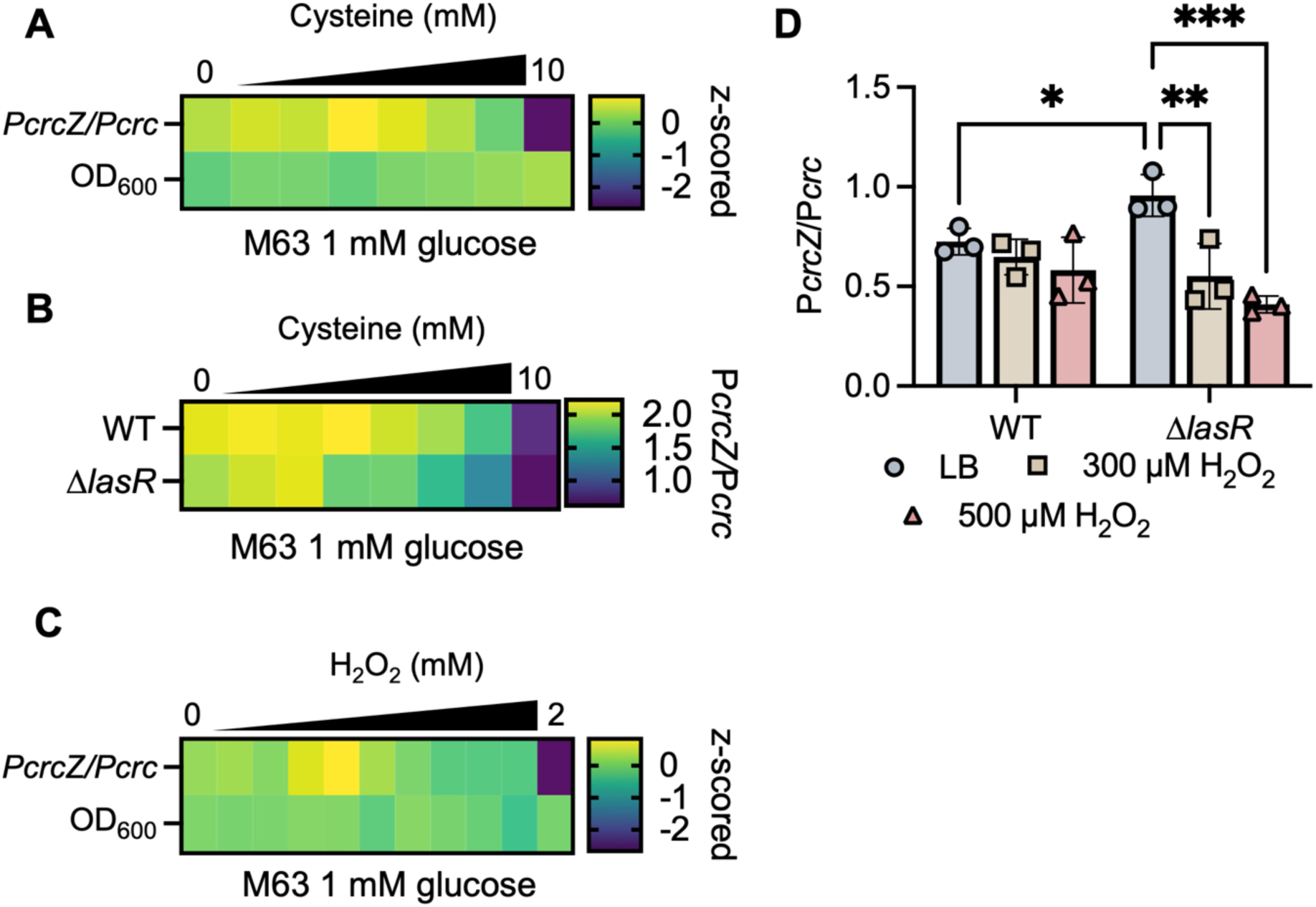
Low levels of reducing or oxidizing agents promoted P*crcZ* activity and high levels repressed P*crcZ* activity. **A)** The PA14 wild type was grown in M63 with 1 mM glucose in a 96-well plate. Cysteine was serially diluted 1:2 from 10 mM to 0.156 mM. P*crcZ* activity and OD600 after 12 h growth were z-scored and shown in a heat map. P*crcZ* activity is highest with the addition of 0.625 mM cysteine and significantly repressed by 10 mM cysteine (p < 0.0001). The OD600 did not change across cysteine concentrations. Statistical analysis was performed on the data prior to z-scoring using a two-way ANOVA and Dunnett’s multiple comparisons test comparing each condition. **B)** The wild type and Δ*lasR* mutant were grown in M63 with 1 mM glucose in a 96-well plate. Cysteine was serially diluted 1:2 from 10 mM to 0.156 mM. Low amounts of cysteine promoted P*crcZ* activity and high amounts repressed P*crcZ* activity in both strains. P*crcZ* activity in the Δ*lasR* mutant was more sensitive to repression by cysteine and significantly repressed by both 5 and 10 mM cysteine (p < 0.0026). P*crcZ* activity in the wild type was only significantly repressed by 10 mM cysteine (p < 0.0001). Statistical analysis was performed using a two-way ANOVA and Sidak’s multiple comparisons test comparing each condition (n = 3). **C)** The PA14 wild type was grown in M63 with 1 mM glucose in a 96-well plate. H2O2 was serially diluted 1:2 from 2 mM to 0.00195 mM (0.125 mM H2O2 is not shown). P*crcZ* activity and OD600 after 12 h growth were z-scored and shown in a heat map. P*crcZ* activity was highest with the addition of 15.6 µM H2O2 and significantly repressed by the addition of 2 mM H2O2 (p < 0.0001). The OD600 did not change across H2O2 concentrations. Statistical analysis was performed on the data prior to z-scoring using a two-way ANOVA and Dunnett’s multiple comparisons test comparing each condition. **D)** *P. aeruginosa* was grown in LB liquid medium with 300 or 500 µM H2O2 for 16. The Δ*lasR* mutant had significantly elevated P*crcZ* activity in LB and this was repressed by the addition of H2O2 such that activity was not significantly different from the wild type. P*crcZ* activity in the wild type was not significantly repressed by H2O2 (p > 0.31). Statistical analysis was performed with a two-way ANOVA Tukey’s multiple comparisons tests comparing strains and conditions (n = 3, * = p < 0.05, ** = p < 0.01, *** = p < 0.001).

As we had observed differences in intracellular cysteine and GSH in the Δ*lasR* mutant relative to the wild type, we proposed that the Δ*lasR* mutant may be more sensitive to P*crcZ* repression by cysteine. As we showed before (**Fig. 4B**), the wild type and Δ*lasR* mutant had similar levels of P*crcZ* activity in M63 with 1 mM glucose. However, we found that the Δ*lasR* mutant was sensitive to P*crcZ* repression by lower amounts of cysteine than the wild type (**Fig. 5B**). P*crcZ* activity in the Δ*lasR* mutant was significantly repressed by 5 or 10 mM cysteine while P*crcZ* activity in the wild type was only significantly repressed by 10 mM cysteine.

Given that P*crcZ* activity is repressed by low and high levels of cysteine, we hypothesized that CbrA-CbrB activity is sensitive to the redox state of the cell. To determine if other modulators of the cellular redox state alter P*crcZ* activity, we tested the effects of H_2_O_2_ on P*crcZ* activity in the wild type in M63 with 1 mM glucose. Small amounts of H_2_O_2_ (15.6 µM) promoted P*crcZ* activity while 2 mM significantly repressed activity (**Fig. 5B**). These levels of H_2_O_2_ did not impede growth.

We found that in M63 with 1 mM glucose, P*crcZ* activity was elevated in both the wild type and Δ*lasR* mutant and similarly sensitive to repression by exogenous cysteine (**Fig. 4C**). However, in LB medium P*crcZ* activity was only elevated and sensitive to cysteine or GSH-mediated repression in the Δ*lasR* mutant (Fig. 3B and **F**). Thus, we hypothesized that in LB medium only the Δ*lasR* mutant would be sensitive to P*crcZ-*repression by H_2_O_2_. We grew the PA14 wild type and Δ*lasR* mutant in subinhibitory (**Fig. S4C**) concentrations of H_2_O_2_ and found that P*crcZ* activity in the Δ*lasR* mutant was significantly repressed to wild-type levels by the addition of H_2_O_2_ and the wild type was unaffected (**Fig. 5D**). Taken together, our results demonstrate that small shifts in the cellular redox environment can promote CbrA-CbrB activity while larger changes repress activity.

### Relief of Crc-mediated repression suppressed P*crcZ* activity

We found that P*crcZ* activity was significantly decreased in the Δ*crc* mutant and restored by complementation (**Fig. 6A**). Crc has been implicated in the regulation of cysteine biosynthesis enzymes (26, 27). However, neither GSH nor cysteine were significantly different in the Δ*crc* mutant or complement grown in LB for 12 h. We found that intracellular GSH was significantly elevated in a Δ*cbrB* mutant and this is rescued by complementation while cysteine levels were not significantly different (**Fig. 6B**). Thus, while CbrB is necessary to maintain the GSH balance, differences in GSH or cysteine levels do not explain P*crcZ* repression in a Δ*crc* mutant.

**Fig. 6.**
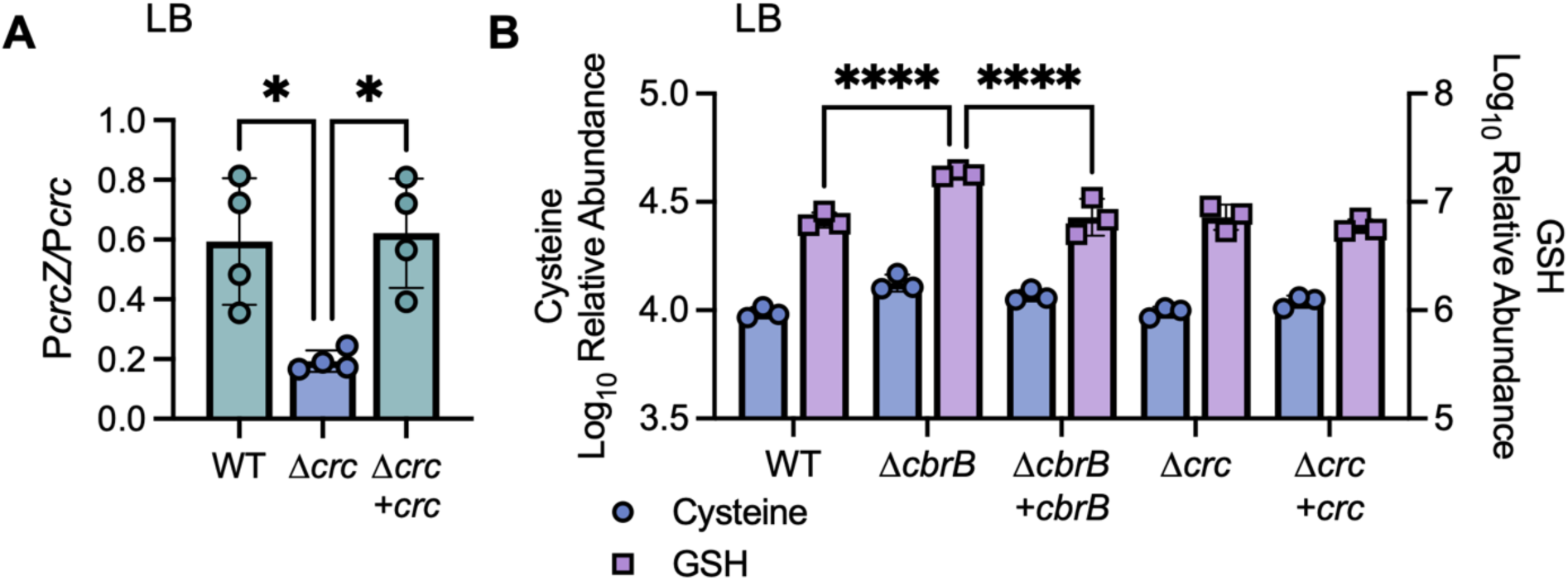
The Δ*crc* mutant had low *PcrcZ* activity. **A)** P*crcZ* activity was significantly lower in the Δ*crc* mutant after 16 h growth in LB relative to the wild type or Δ*crc+crc* complement. Statistical analysis was performed with an ordinary one-way ANOVA and Tukey’s multiple comparisons tests comparing each strain (n = 3). **B)** Cysteine levels were not different between the wild type, Δ*cbrB* mutant, Δ*crc* mutant, or complements (p > 0.21) after 12 h growth in LB. GSH levels were significantly higher in the Δ*cbrB* mutant relative to the wild type or Δ*cbrB+cbrB* complement but unchanged in the Δ*crc* mutant or complement (p > 0.65). The wild type data shown here is also shown in Fig. 2E and Fig. S2E. Statistical analysis was performed with a two-way ANOVA and Tukey’s multiple comparisons tests comparing each strain. For all panels * = p < 0.05, ** = p < 0.01, *** = p < 0.001, and **** = p < 0.0001.

### Stationary phase signaling affects P*crcZ* activity with and without corresponding changes in thiol pools

There are many regulators with increased activity in stationary phase cultures when P*crcZ* activity is elevated (**Fig. 2C**). We previously found that the other quorum sensing regulators RhlR and PqsR, as well as redox-active phenazines under RhlR control, can repress CbrB-dependent phosphatase activity (8). Here we found that the absence of RhlR or PqsR did not affect P*crcZ* activity (**Fig. S5A**). However, the phenazine-null Δ*phzA1-G1ΔphzA2-G2* mutant (referred to as ΔΔ*phz*) did have significantly elevated P*crcZ* activity relative to the wild type (**Fig. S5B**). Phenazines can be elevated in a Δ*crc* mutant (28) but loss of phenazines did not restore P*crcZ* activity in a Δ*crc* background (**Fig. S5C**).

The alarmone ppGpp accumulates in late-stage cultures and has been implicated in repressing cysteine biosynthesis (29–31). Deletion of the two ppGpp synthases, RelA and SpoT, significantly increased P*crcZ* activity relative to the wild type in LB liquid medium (**Fig. S5D**). However, we observed no significant change in cysteine or GSH levels in the Δ*relAΔspoT* mutant compared to the wild type (**Fig. S5E**). Loss of ppGpp had the opposite effect in a Δ*lasR* mutant, significantly repressing P*crcZ* activity (**Fig. S5F**). The Δ*lasRΔrelAΔspoT* mutant had significantly higher intracellular GSH but no change in cysteine levels relative to the Δ*lasR* mutant (**Fig. S5G**). This was consistent with our previous findings that GSH repressed P*crcZ* in a Δ*lasR* mutant (**Fig. 3F**). The effects of ppGpp on P*crcZ* activity and GSH levels are pleiotropic and influenced by the LasR status of the cell.

Evans, *et al.* found that the sigma factor RpoS represses P*crcZ* activity (18). The Δ*rpoS* mutant had significantly elevated P*crcZ* activity in LB and this was rescued by complementation of *rpoS* at the native locus (**Fig. S5H**). However, there were no significant difference in cysteine or GSH levels between the wild type, Δ*rpoS* mutant, or complement (**Fig. S5I**). Many stationary phase factors including phenazines, ppGpp, and RpoS altered P*crcZ* activity. Some of these effects correlated with changes in GSH but others did not.

### Cysteine repressed Δ*lasR* mutant competitive fitness

We previously proposed the model that elevated CbrA-CbrB activity in *lasR* mutants provides a competitive fitness advantage (10). Here, we found that coculture with the wild type increased P*crcZ* activity in the Δ*lasR* mutant. We incubated a Δ*lasR* mutant with the P*crcZ*P*crc* dual promoter fusion (30%) with an unlabeled Δ*lasR* or wild type (70%) for 12 h in LB. The Δ*lasR* mutant with the promoter fusion had significantly more P*crcZ* activity (normalized to P*crc*) when cocultured with the wild type (**Fig. 7A**). We then tested competitive fitness by co-inoculating a labeled wild-type strain (70%) with an unlabeled competitor (30%) and incubating the cultures for 12 h in LB. The unlabeled wild type had no competitive advantage over the labeled strain and the Δ*lasR* mutant is more fit than the wild type (**Fig. 7B**). Consistent with our model, the Δ*lasRΔcbrA* mutant was significantly less fit than the Δ*lasR* mutant in competitions with the wild type (**Fig. 7C**).

**Fig. 7.**
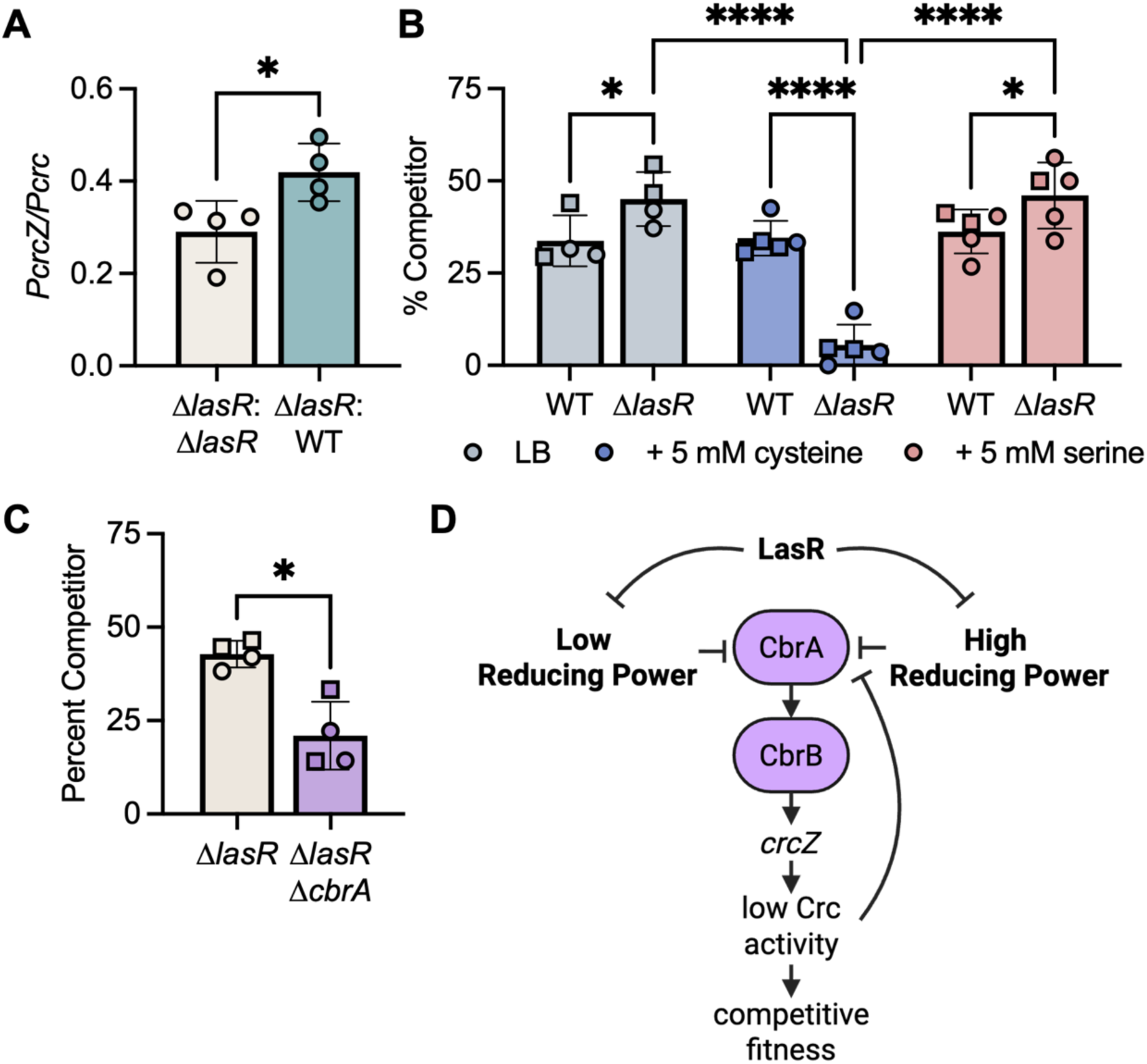
CbrA activity was required for the Δ*lasR* mutant competitive fitness. All data shown in Fig. 7 was collected after 12 h growth in LB medium. **A)** P*crcZ* activity was significantly elevated in the Δ*lasR* mutant when incubated with unlabeled wild type (30:70) relative to when incubated with an unlabeled Δ*lasR* mutant. Statistical analysis was performed with a two-way unpaired Welch’s t-test. **B)** Wild type and Δ*lasR* were competed against a labeled wild-type strain (30:70) for 12 h. The Δ*lasR* mutant outcompeted the wild type in LB and with serine but was significantly less fit with cysteine. Statistical analysis was performed with a two-way ANOVA and Tukey’s multiple comparisons tests comparing each strain and condition. **C)** The Δ*lasR* and Δ*lasRΔcbrA* mutants were competed against a labeled wild-type strain (30:70). The Δ*lasR* mutant is significantly more fit than the Δ*lasRΔcbrA* mutant. Statistical analysis was performed with a two-way unpaired Welch’s t-test. For all panels * = p < 0.05, ** = p < 0.01, *** = p < 0.001, and **** = p < 0.0001. **D)** A model of LasR regulation of CbrA activity. We propose LasR regulates the reducing power in the cell. Both low and high reducing power repress CbrA activity, leading to decreased competitive fitness.

Given our findings that exogenous cysteine repressed P*crcZ* activity, we tested the effects of cysteine on the Δ*lasR* mutant competitive fitness. The addition 5 mM cysteine did not affect the wild type fitness but significantly decreased the Δ*lasR* mutant competitive fitness while the addition of 5 mM serine had no effect. In single strain cultures, the Δ*lasR* mutant did not show a growth defect in LB with 5 mM cysteine (**Fig. S3A**). These data support our model that elevated CbrA-CbrB activity in the Δ*lasR* mutant promotes competitive fitness, and this is sensitive to the redox state of the cell (see **Fig. 7D** for model).

## Discussion

Loss-of-function (LOF) mutations to the gene encoding the quorum sensing regulator LasR are commonly observed in both clinical and environmental *Pseudomonas aeruginosa* isolates (2, 4) and are associated with worse clinical outcomes (1, 7). The worse outcomes associated with *lasR* mutants may seem paradoxical as active quorum sensing is associated with increased virulence. We and others have demonstrated that another quorum sensing regulator, RhlR, can be activated even the absence of LasR (9, 32, 33). Yet the rescue of RhlR function does not explain the repeated emergence or worse outcomes associated with *lasR* mutants. We previously found that the two-component system CbrA-CbrB is necessary for the evolution of *lasR* mutants (10) and increased virulence gene expression, including *rhlR*, in physiologically relevant conditions (8). CbrA-CbrB is specific to and highly conserved within *Pseudomonas spp.* and known for its translational regulation of carbon catabolism (11, 13, 15) as well as virulence (16, 17, 34). However, the mechanism of activation for CbrA has not been established. Here we present a model wherein CbrA-CbrB activity requires moderate reducing power and is sensitive to changes in the cellular thiol pool.

Together our data strongly suggest a model in which intracellular thiols must be within a specific range to allow for CbrA-CbrB activity. We validated our previous findings that CbrA-CbrB activity is elevated in *lasR* mutants (8, 10) (**Fig. 1B-C, Fig. S1F**). Then, leveraging CbrA-CbrB-dependent phosphatase activity, we developed a high-throughput screen to identify genes necessary for elevated CbrA-CbrB activity in the LasR-J215 isolate. We found that disrupting genes in the cysteine and glutathione (GSH) biosynthesis pathway abolished CbrA-CbrB-dependent phosphatase (**Fig. 1E-I**, **Fig. 1K-L**) and supplementation with exogenous cysteine rescued phosphatase activity in these mutants (**Fig. 1G-I**). Furthermore, cysteine levels elevated in the Δ*lasR* mutant prior to a significant increase in P*crcZ* activity (**Fig. 2C-D**). Unexpectedly, we found that high levels of exogenous cysteine or GSH repressed P*crcZ* activity in the Δ*lasR* mutant (**Fig. 3A-C, 3F**). However, small amounts of cysteine stimulated P*crcZ* activity in both the Δ*lasR* mutant and wild type (**Fig. 5B**). We observed the same pattern with the oxidizing agent H_2_O_2_, where small amounts promoted P*crcZ* activity and high amounts repressed it (**Fig. 5C-D**). Thus, we propose that CbrA-CbrB activity is sensitive to the redox state of the cell and repressed by both low and high levels of reducing power.

The observation that cysteine represses P*crcZ* activity in stationary phase cultures (**Fig. 2C, 4D**) suggests a connection between these cues (**Fig. 4A-B**). We investigated the effects of stationary phase signaling on P*crcZ* activity and cysteine and GSH abundance. We found that cysteine increases in stationary phase in the Δ*lasR* mutant in LB (**Fig. 2D**). Previous work has implicated the stationary phase alarmone ppGpp in repressing cysteine biosynthesis (29, 30). We did observe a significant increase in GSH abundance in the ppGpp-null Δ*lasRΔrelAΔspoT* mutant relative to the Δ*lasR* mutant (**Fig. S5G**). As expected, given our findings that high levels of GSH repress P*crcZ* activity (**Fig. 3B, E-F**), this correlated with a significant decrease in P*crcZ* activity (**Fig. S5F**). However, in the LasR+ background we found no significant change in either cysteine or GSH levels in the Δ*relAΔspoT* mutant (**Fig. S5E**). Loss of ppGpp did significantly increase P*crcZ* activity in the PA14 LasR+ background (**Fig. S5D**). Thus, ppGpp represses CbrA-CbrB activity in a LasR+ strain and stimulates CbrA-CbrB activity in a LasR-strain. GSH may be a critical factor in a *lasR* mutant but it is unclear how ppGpp represses activity in a LasR+ strain. In a similar vein, we validated the previous findings that the sigma factor RpoS represses P*crcZ* activity (18) but observed no change in cysteine or GSH levels in this mutant (**Fig. S5H-I**). It should be noted that our analysis of the thiol pool was limited to two of several thiol-containing metabolites that can modulate the redox state of the cell. Recent work has demonstrated that fluorescein-based dye assays are not a reliable method for investigating the redox state of a cell (35). Further analysis of other thiol-containing metabolites like homocysteine or quantifying the NADH/NAD+ ratio may provide further insight into how stationary phase-associated factors influence P*crcZ* activity.

We also found that phenazines repress P*crcZ* activity (**Fig. S5B**). Phenazines are redox-active small molecules that induce oxidative stress in host cells (36) but play an important role in maintaining the *Pseudomonas* redox state. Loss of phenazines leads to an increase in cellular reducing power as evidenced by an increase in the NADH/NAD+ ratio and rate of pyocyanin reduction (37, 38). We propose that late-stage phenazine production in the wild type is one contributing factor repressing the reducing power and subsequently CbrA-CbrB activity. *lasR* mutants generally produce fewer phenazines (39) and this may contribute to their elevated CbrA-CbrB activity. However, we have shown that in coculture with LasR+ cells, the Δ*lasR* mutant produces more phenazines (40) and yet P*crcZ* activity in the *ΔlasR* mutant is elevated in coculture with the wild type (**Fig. 7A**). Further research is needed to elucidate if phenazines impact CbrA-CbrB activity and subsequently competitive fitness in heterogenous populations.

Interestingly, loss of Crc, the downstream repressor in the CbrA-CbrB pathway, resulted in significantly less P*crcZ* activity (**Fig. 6A**). Crc has been implicated in regulating cysteine biosynthesis (26, 41), however we found no significant differences in cysteine or GSH levels in the Δ*crc* mutant in these conditions (**Fig. 6B**). We did find significantly more GSH in the Δ*cbrB* mutant relative to the wild type or complement. Taken together with our screening results, we propose that moderate levels of GSH inside the cell promote CbrA-CbrB activity and in the absence of CbrB, GSH continues to accumulate. This data aligns with our previous findings that deletion of *cbrB* increases methylglyoxal resistance in a Δ*lasR* mutant (25). We propose that CbrA-CbrB activity may repress thiol pools in *P. aeruginosa* and so alter sensitivity to oxidative stresses. We have only found Crc LOF mutants arising in the absence of CbrA or CbrB (10). This suggests *crc* mutants are less fit and thus feedback inhibition to prevent overactivation of CbrA-CbrB may contribute to fitness. Further analysis of other redox-modulating factors and possible Crc-mediated regulation will provide more insight into feedback inhibition of CbrA-CbrB activity and CbrA-CbrB-dependent stress sensitivity.

Throughout this study we found multiple lines of evidence suggesting *lasR* mutants regulate intracellular thiol pools differently from LasR+ cells. In some contexts (MOPS-glucose agar or liquid LB), the *lasR* mutants had significantly more intracellular cysteine than the LasR+ comparator strain (**Fig. 2B, D**). Previously we reported that *lasR* mutants grown on LB agar have significantly less intracellular cysteine and GSH than the wild type (9, 25). Supplementing exogenous cysteine to the media significantly increased cysteine in the wild type (**Fig. 3D**) but significantly increased GSH in the Δ*lasR* mutant (**Fig. 3E**). Previous work has implicated LasR directly in the regulation of *cysI* (42). Cysteine biosynthesis and accumulation is tightly regulated at multiple levels as cysteine is necessary to protect cells from oxidative stress but can also become toxic at high levels by promoting Fenton chemistry and subsequent production of hydroxyl radicals. Thus, cysteine is converted to the less toxic GSH or secreted from cells the prevent toxicity (43–46). Our results indicate that LasR regulates the biosynthesis and/or accumulation of thiol-containing metabolites, but the mechanisms of that regulation are not yet fully established.

We propose these differences in thiol pool regulation contribute to the planktonic lifestyle associated with *lasR* mutants. *las* mutants are typically poor biofilm formers (47) and LasR plays a less critical role in regulating the transcriptome in biofilms than planktonic cultures (42). We found that the Δ*lasR* mutant had less P*crcZ* activity compared to the wild type on LB agar (**Fig. S2C**), unlike in LB liquid where it had more (**Fig. 3B**), and these differences correlated with intracellular cysteine abundance. Biofilms protect cells from stressors like antibiotics (48) and so increased cysteine production may be an aspect of Δ*lasR* mutants adapting to a planktonic lifestyle where they are more exposed to oxidative stressors. The subsequent increase in CbrA-CbrB activity provides a growth and competitive advantage to the Δ*lasR* mutants in planktonic conditions (**Fig. 7B**) (9, 10). Interestingly, we found that exogenous cysteine not only repressed the competitive fitness advantage of the Δ*lasR* mutant but sensitized it to competition by the wild type (**Fig. 7B**). Counter to previously held models, Δ*lasR* mutants are not sensitive to cyanide policing by the wild type (49). We propose that CbrA-CbrB activity may play a role in protecting Δ*lasR* mutants from cyanide or similar antagonistic factors produced by other microbes or host cells. Others have reported that *cbrA* or *cbrB* mutants are more resistant to antibiotics and killing or uptake by immune cells (16, 34). Thus, we propose that under high levels of redox stress, it is more favorable for cells to repress CbrA-CbrB activity. CbrA-CbrB inactivation at both high and low levels of reducing power enables *P. aeruginosa* to survive in ever-changing environments.

The dual layer of transcriptional and translational regulation enabled by CbrA-CbrB activity allows *Pseudomonas* to quickly express condition-dependent proteins. The mRNA transcripts under Crc control are also under unique transcriptional regulation. This transcriptional regulation allows for specific, but slower responses. Expression of *crcZ* leads to rapid, universal relief of Crc-mediated translational repression. Changes in the cellular redox state are a near universal signal of stress or changes in metabolic capacity and thus an elegant mechanism for activating CbrA-CbrB across diverse conditions.

## Materials and Methods

### Strains and growth conditions

Bacterial strains and plasmids used in this study are listed in Table S2. Bacteria were streaked from frozen stocks onto lysogeny broth (LB) with 1.5% agar (50). Planktonic cultures were grown in 5 mL LB medium in 18 mm borosilicate glass tubes on a roller drum at 37°C.

### Construction of in-frame deletion and complementation plasmids

Construction of in-frame deletion and complementation constructs was performed using yeast cloning techniques in *Saccharomyces cerevisiae* as previously described (51) or with the NEBuilder HiFi DNA Assembly Cloning Kit (NEB #E5520S). Constructs for in-frame deletion of most or all the coding sequence and chromosomal complementation were made using the allelic replacement vector pMQ30 with 1 kb homology on either side of the region to be deleted or complemented. For all complements, the gene under its native promoter was reconstituted. Plasmids were purified from yeast using Zymoprep Yeast Plasmid Miniprep II (Zymo #D2004) according to the manufacturer’s protocol and transformed into the *Escherichia coli* strain S17/λpir by electroporation. Plasmids were introduced into *P. aeruginosa* by conjugation, and recombinants were obtained using sucrose counter selection. Genotypes were screened by PCR, and plasmid constructs were confirmed by sequencing.

### J215 genome assembly

PAO1 and PA14 genomic sequences and features were accessed from the latest Genbank genomes on PGD (GCF_000006765.1 and GCF_000014625.1). The J215 genome was constructed via hybrid assembly and annotated using SPAdes and Prokka. J215 predictive features were annotated by sequence identity using alignments against the PAO1 and PA14 reference genomes. Annotated genes that aligned exactly end-to-end were annotated as exact matches. All remaining J215 genes and PAO1 genes that were unmatched were extracted and realigned against each other. One-to-one partial matches were annotated with the PAO1 feature information. Any remaining unmatched J215 predictive features were then run through the same process against the PA14 genome. Any remaining J215 predictive features were annotated as such.

### Screen of J215 clinical isolate transposon mutant library

The J215 clinical isolate was randomly mutagenized with a Mariner Transposon and ∼7,500 individual isolates stocked in 96-well plates. These isolates were grown in 96-well plates in LB for 16 hours then plated on MOPS-glucose agar with 1 mM Pi and BCIP using a frogger. Isolates with BCIP phenotypes were tested again in triplicate with 5 µL spots of OD_600_ = 1 culture as described above.

Arbitrary PCR to amplify the genetic sequence after the transposon and Sanger sequencing via the Dartmouth Genomics Shared Resource core facility was used to identify the approximate transposon insertion location in the J215 genome.

### Construction of dual promoter fusion

The promoter region of *crcZ* (161bp) was amplified from PA14 gDNA and the mOrange2 coding sequence was amplified from pUC18miniTn7::Pa1104103-mOrange2, each with homologous overlap. The two fragments were inserted upstream of the P*crc* promoter region in the pSEK109.P*crc_mScarlet* plasmid via the NEBuilder HiFi DNA Assembly Cloning Kit (NEB #E5520S) then transformed into DH5*a E. coli*. Successful construction was confirmed with sequencing, and the construct was transformed into the *E. coli* strain S17/λpir by electroporation and then introduced into *P. aeruginosa* by conjugation. The gentamycin resistance cassette can be flipped out but was not removed in these strains.

### P*crcZ* activity assays and microscopy

*P. aeruginosa* strains containing the P*crcZ*P*crc* dual promoter fusion integrated at the *attB* site were grown overnight as described above. Cultures were then normalized to OD_600_ = 1 in LB and either plated as 5 µL spots on agar or inoculated in 5 mL tubes or 96 well plates at a starting density of OD_600_ = 0.05 and incubated at 37°C, shaking for liquid cultures. To quantify P*crcZ* activity, colony biofilms were removed from agar using a pipette tip and resuspended in 100 µL water in a clear bottom, black 96-well plate. Liquid cultures were transferred to 96 well plates. The plates were shaken for 5 seconds and OD_600_ and RFU at excitation 545/10 emission 568/10 (P*crcZ*_*mOrange2*) and excitation 569/10 and emission 594/10 (P*crc*_*mScarlet*) were read in a Synergy Neo2 multi-mode plate reader, blanked to water or media only controls. P*crcZ* RFU was normalized to P*crc* RFU.

Confocal microscopy images of promoter fusion strains were taken by spotting 3 µL of bacterial culture on 1% agarose pads to immobilize live cells. Confocal images were acquired with a Nikon AX laser scanning confocal system, fitted with an Apo 40x/1.25 NA water objective running Nikon NIS-Elements (AR 5.41.02.). A 561-nm laser line was used to excite the mOrange2 (emission at 571-593-nm) driven by the *crcZ* promoter and the mScarlet (emission at 601-651-nm) driven by the *crc* promoter. For single cell analysis each sample was spotted onto three agarose pads, and three images were taken of each pad (total of nine images per sample). Images were thresholded and intensity measurements per area were quantified using Nikon NIS-Elements using Fill Area (µm 2), Mean Intensity, and Count features. Analysis was automated using General Analysis. Objects with a fill area greater than 4 µM or smaller than 0.7 µM were excluded from analysis. The mOrange2 Mean Intensity of each object was normalized to the mScarlet Mean Intensity to calculate P*crcZ/*P*crc* activity for individual cells.

### Phosphatase activity assessment with BCIP

Agar plates were made as previously described (8, 21) with MOPS minimal medium (52) with 20 mM glucose, 15 g/L agar (referred to here as MOPS-glucose agar), 1 mM K_2_HPO_4_/KH_2_PO_4_, and 60 µg/mL BCIP (Sigma Aldrich #11383221001). Overnight cultures of *P. aeruginosa* were grown in LB, normalized to an OD_600_ = 1 in LB, and then 5 µL was spotted on MOPS-glucose agar and incubated at 37°C for 16 hours.

### Cysteine and GSH quantification

*P. aeruginosa* was grown overnight as described above. Cultures were then normalized to OD_600_ = 1 in LB and either plated as 5 µL spots in triplicate-quintuplicate on agar or inoculated in 5 mL tubes at a starting density of OD_600_ = 0.05 and incubated at 37°C for 12 h. The colony biofilms for each sample were collected together from the agar using a pipette tip. For liquid cultures, 1-2 mL of culture was pelleted and the supernatant removed. Samples were then snap frozen in microcentrifuge tubes with 200 proof ethanol and dry ice and stored at −80°C until all biological replicates were collected.

Metabolites were extracted by resuspending the cells in 0.2 – 1 mL (depending on pellet size) −80°C 80% methanol, vortexing the samples, and incubating at −20°C for 5 - 10 minutes for two cycles. To ensure complete cell lysis, samples were sonicated three times at 10-30% amplitude (depending on sample volume) for 5 seconds while on ice. Lysate was incubated at −20°C overnight to precipitate proteins then vortexed once more and centrifuged at 20,000 x g for 15 min at 4°C. The protein pellet was dissolved in 3M urea at 65°C and total protein was quantified via Bradford assay. The supernatant was transferred to a new microcentrifuge tube and metabolites were dried to a pellet via SpeedVac and stored at −80°C. Metabolite pellets were shipped to the Northwestern University Metabolomics Core Facility and semi-quantitative analysis of cysteine and GSH (injection volume normalized to total protein) was determined via liquid chromatography – mass spectrometry (LC-MS).

### Competition assays

70% labeled wild type *P. aeruginosa* constitutively expressing *lacZ* was inoculated with 30% unlabeled competitor strain to a final starting OD_600_ = 0.05 in 5 mL LB liquid. Cultures were incubated on a roller drum for 12 h then serially diluted. 10^-5^ – 10^-7^ dilutions of cultures were plated on LB agar with 150 µg/mL 5-Bromo-4-Chloro-3-Indolyl β-D-Galactopyranoside (X-gal). The *lacZ*-expressing strains convert X-gal to a blue pigment after overnight incubation. The percentage of white (competitor) colonies was calculated.

### Statistical analysis and figure design

Statistical analyses were performed and figures made with GraphPad Prism version 11.1 for macOS. **Fig. 1A, S1A, 2A**, and **7D** were created in BioRender (y62m090).

## Acknowledgements

This work was made possible by support from the NIH (ACF31AI179113, LMF31AI191725, NHLBI T32HL134598 (AC), T32AI007519 (LM), and R01HL169973 (JB)), Cystic Fibrosis Foundation (BOMBER24G0), Dartmouth BioMT core (NIH NIGMS P20-GM113132 and P30-GM165328), and the Metabolomics Core Facility at Robert H. Lurie Comprehensive Cancer Center (generously supported by NCI CCSG P30 CA060553) of Northwestern University.

## References

1. Hoffman LR, Kulasekara HD, Emerson J, Houston LS, Burns JL, Ramsey BW, Miller SI. 2009. *Pseudomonas aeruginosa lasR* mutants are associated with cystic fibrosis lung disease progression. Journal of Cystic Fibrosis 8:66–70.

2. Groleau MC, Taillefer H, Vincent AT, Constant P, Déziel E. 2022. *Pseudomonas aeruginosa* isolates defective in function of the LasR quorum sensing regulator are frequent in diverse environmental niches. Environmental Microbiology 24:1062–1075.

3. O’Connor K, Zhao CY, Mei M, Diggle SP. 2022. Frequency of quorum-sensing mutations in *Pseudomonas aeruginosa* strains isolated from different environments. Microbiology 168:001265.

4. Trottier MC, de Oliveira Pereira T, Groleau M-C, Hoffman LR, Dandekar AA, Déziel E. 2024. The end of the reign of a “master regulator’’? A defect in function of the LasR quorum sensing regulator is a common feature of *Pseudomonas aeruginosa* isolates. MBio 15:e02376–23.

5. Hentzer M, Wu H, Andersen JB, Riedel K, Rasmussen TB, Bagge N, Kumar N, Schembri MA, Song Z, Kristoffersen P. 2003. Attenuation of *Pseudomonas aeruginosa* virulence by quorum sensing inhibitors. The EMBO journal 22:3803–3815.

6. Antunes LCM, Ferreira RB, Buckner MM, Finlay BB. 2010. Ǫuorum sensing in bacterial virulence. Microbiology 156:2271–2282.

7. Hammond JH, Hebert WP, Naimie A, Ray K, Van Gelder RD, DiGiandomenico A, Lalitha P, Srinivasan M, Acharya NR, Lietman T. 2016. Environmentally endemic *Pseudomonas aeruginosa* strains with mutations in lasR are associated with increased disease severity in corneal ulcers. MSphere 1:10.1128/msphere.00140-16.

8. Conaway A, Mould DL, Todorovic I, Hogan DA. 2025. Loss of LasR function leads to decreased repression of *Pseudomonas aeruginosa* PhoB activity at physiological phosphate concentrations. Journal of Bacteriology 207:e00189–24.

9. Mould DL, Finger CE, Conaway A, Botelho N, Stuut SE, Hogan DA. 2024. Citrate cross-feeding by *Pseudomonas aeruginosa* supports *lasR* mutant fitness. mBio 15:e01278–23.

10. Mould DL, Stevanovic M, Ashare A, Schultz D, Hogan DA. 2022. Metabolic basis for the evolution of a common pathogenic *Pseudomonas aeruginosa* variant. Elife 11:e76555.

11. Sonnleitner E, Abdou L, Haas D. 2009. Small RNA as global regulator of carbon catabolite repression in *Pseudomonas aeruginosa*. Proceedings of the National Academy of Sciences 106:21866–21871.

12. Abdou L, Chou H-T, Haas D, Lu C-D. 2011. Promoter Recognition and Activation by the Global Response Regulator CbrB in *Pseudomonas aeruginosa*. Journal of Bacteriology 193:2784–2792.

13. Nishijyo T, Haas D, Itoh Y. 2001. The CbrA–CbrB two-component regulatory system controls the utilization of multiple carbon and nitrogen sources in *Pseudomonas aeruginosa*. Molecular microbiology 40:917–931.

14. García-Mauriño SM, Pérez-Martínez I, Amador CI, Canosa I, Santero E. 2013. Transcriptional activation of the CrcZ and CrcY regulatory RNAs by the CbrB response regulator in *Pseudomonas putida*. Molecular Microbiology 89:189–205.

15. Valentini M, García-Mauriño SM, Pérez-Martínez I, Santero E, Canosa I, Lapouge K. 2014. Hierarchical management of carbon sources is regulated similarly by the CbrA/B systems in *Pseudomonas aeruginosa* and *Pseudomonas putida*. Microbiology 160:2243–2252.

16. Yeung AT, Janot L, Pena OM, Neidig A, Kukavica-Ibrulj I, Hilchie A, Levesque RC, Overhage J, Hancock RE. 2014. Requirement of the *Pseudomonas aeruginosa* CbrA sensor kinase for full virulence in a murine acute lung infection model. Infect Immun 82:1256–67.

17. Gil-Gil T, Valverde JR, Martínez JL, Corona F. 2023. In vivo genetic analysis of *Pseudomonas aeruginosa* carbon catabolic repression through the study of CrcZ pseudo-revertants shows that Crc-mediated metabolic robustness is needed for proficient bacterial virulence and antibiotic resistance. Microbiol Spectr 11:e0235023.

18. Evans CR, Smiley MK, Asahara Thio S, Wei M, Florek LC, Dayton H, Price-Whelan A, Min W, Dietrich LEP. 2023. Spatial heterogeneity in biofilm metabolism elicited by local control of phenazine methylation. Proceedings of the National Academy of Sciences 120:e2313208120.

19. Hammond JH, Dolben EF, Smith TJ, Bhuju S, Hogan DA. 2015. Links between Anr and quorum sensing in *Pseudomonas aeruginosa* biofilms. Journal of bacteriology 197:2810–2820.

20. Horwitz JP, Chua J, Noel M, Donatti JT, Freisler J. 1966. Substrates for cytochemical demonstration of enzyme activity. II. Some Dihalo-3-indolyl Phosphates and Sulfates1. Journal of medicinal chemistry 9:447–447.

21. Doing G, Koeppen K, Occipinti P, Harty CE, Hogan DA. 2020. Conditional antagonism in co-cultures of *Pseudomonas aeruginosa* and *Candida albicans*: an intersection of ethanol and phosphate signaling distilled from dual-seq transcriptomics. PLoS genetics 16:e1008783.

22. Wanner BL. 1996. Phosphorus assimilation and control of the phosphate regulon. Escherichia coli and Salmonella: cellular and molecular biology, 2nd ed ASM Press, Washington, DC 41:1357–1381.

23. Gardner SG, Johns KD, Tanner R, McCleary WR. 2014. The PhoU protein from *Escherichia coli* interacts with PhoR, PstB, and metals to form a phosphate-signaling complex at the membrane. Journal of bacteriology 196:1741–1752.

24. Liberati NT, Urbach JM, Miyata S, Lee DG, Drenkard E, Wu G, Villanueva J, Wei T, Ausubel FM. 2006. An ordered, nonredundant library of *Pseudomonas aeruginosa* strain PA14 transposon insertion mutants. Proc Natl Acad Sci U S A 103:2833–8.

25. Altamirano A, Ruzic M, Cryan AE, Hogan DA. 2025. Effects of low intracellular glutathione and CbrA-CbrB-Crc signaling on methylglyoxal sensitivity in *Pseudomonas aeruginosa* LasR-deficient mutants. Journal of Bacteriology 207:e00394–24.

26. Corona F, Reales-Calderón JA, Gil C, Martínez JL. 2018. The development of a new parameter for tracking post-transcriptional regulation allows the detailed map of the *Pseudomonas aeruginosa* Crc regulon. Scientific Reports 8:16793.

27. Linares JF, Moreno R, Fajardo A, Martínez-Solano L, Escalante R, Rojo F, Martínez JL. 2010. The global regulator Crc modulates metabolism, susceptibility to antibiotics and virulence in *Pseudomonas aeruginosa*. Environ Microbiol 12:3196–212.

28. Huang J, Sonnleitner E, Ren B, Xu Y, Haas D. 2012. Catabolite Repression Control of Pyocyanin Biosynthesis at an Intersection of Primary and Secondary Metabolism in *Pseudomonas aeruginosa*. Applied and Environmental Microbiology 78:5016–5020.

29. Smirnova GV, Tyulenev AV, Bezmaternykh KV, Muzyka NG, Ushakov VY, Oktyabrsky ON. 2019. Cysteine homeostasis under inhibition of protein synthesis in *Escherichia coli* cells. Amino Acids 51:1577–1592.

30. Wu L, Wang Z, Guan Y, Huang X, Shi H, Liu Y, Zhang X. 2020. The (p)ppGpp-mediated stringent response regulatory system globally inhibits primary metabolism and activates secondary metabolism in *Pseudomonas protegens* H78. Applied Microbiology and Biotechnology 104:3061–3079.

31. Dalebroux ZD, Swanson MS. 2012. ppGpp: magic beyond RNA polymerase. Nature Reviews Microbiology 10:203–212.

32. Jensen V, Löns D, Zaoui C, Bredenbruch F, Meissner A, Dieterich G, Münch R, Häussler S. 2006. RhlR expression in *Pseudomonas aeruginosa* is modulated by the *Pseudomonas* quinolone signal via PhoB-dependent and-independent pathways. Journal of bacteriology 188:8601–8606.

33. Soto-Aceves MP, Cocotl-Yañez M, Servín-González L, Soberón-Chávez G. 2021. The Rhl quorum-sensing system is at the top of the regulatory hierarchy under phosphate-limiting conditions in *Pseudomonas aeruginosa* PAO1. Journal of Bacteriology 203:10.1128/jb.00475-20.

34. Yeung AT, Bains M, Hancock RE. 2011. The sensor kinase CbrA is a global regulator that modulates metabolism, virulence, and antibiotic resistance in *Pseudomonas aeruginosa*. J Bacteriol 193:918–31.

35. Korshunov S, Imlay JA. 2025. Fluorescein-based dyes are not valid reporters of oxidative stress in bacteria, and conclusions based on their use must be reconsidered. Proceedings of the National Academy of Sciences 122:e2516113122.

36. Muller M. 2002. Pyocyanin induces oxidative stress in human endothelial cells and modulates the glutathione redox cycle. Free Radic Biol Med 33:1527–33.

37. Price-Whelan A, Dietrich LE, Newman DK. 2007. Pyocyanin alters redox homeostasis and carbon flux through central metabolic pathways in *Pseudomonas aeruginosa* PA14. J Bacteriol 189:6372–81.

38. Ciemniecki JA, Newman DK. 2023. NADH dehydrogenases are the predominant phenazine reductases in the electron transport chain of *Pseudomonas aeruginosa*. Molecular Microbiology 119:560–573.

39. Higgins S, Heeb S, Rampioni G, Fletcher MP, Williams P, Cámara M. 2018. Differential regulation of the phenazine biosynthetic operons by quorum sensing in *Pseudomonas aeruginosa* PAO1-N. Frontiers in cellular and infection microbiology 8:252.

40. Mould DL, Botelho NJ, Hogan DA. 2020. Intraspecies signaling between common variants of *Pseudomonas aeruginosa* increases production of quorum-sensing-controlled virulence factors. mBio 11:e01865–20.

41. Corona F, Martínez JL, Nikel PI. 2019. The global regulator Crc orchestrates the metabolic robustness underlying oxidative stress resistance in *Pseudomonas aeruginosa*. Environmental Microbiology 21:898–912.

42. Jeske A, Arce-Rodriguez A, Thöming JG, Tomasch J, Häussler S. 2022. Evolution of biofilm-adapted gene expression profiles in lasR-deficient clinical *Pseudomonas aeruginosa* isolates. npj Biofilms and Microbiomes 8:6.

43. Park S, Imlay James A. 2003. High Levels of Intracellular Cysteine Promote Oxidative DNA Damage by Driving the Fenton Reaction. Journal of Bacteriology 185:1942–1950.

44. Chonoles Imlay KR, Korshunov S, Imlay JA. 2015. Physiological Roles and Adverse Effects of the Two Cystine Importers of *Escherichia coli*. J Bacteriol 197:3629–44.

45. Ohtsu I, Wiriyathanawudhiwong N, Morigasaki S, Nakatani T, Kadokura H, Takagi H. 2010. The L-Cysteine/L-Cystine Shuttle System Provides Reducing Equivalents to the Periplasm in *Escherichia coli*. Journal of Biological Chemistry 285:17479–17487.

46. Takumi K, Nonaka G. 2016. Bacterial Cysteine-Inducible Cysteine Resistance Systems. J Bacteriol 198:1384–92.

47. Davies DG, Parsek MR, Pearson JP, Iglewski BH, Costerton JW, Greenberg EP. 1998. The Involvement of Cell-to-Cell Signals in the Development of a Bacterial Biofilm. Science 280:295–298.

48. Thöming JG, Häussler S. 2022. *Pseudomonas aeruginosa* Is More Tolerant Under Biofilm Than Under Planktonic Growth Conditions: A Multi-Isolate Survey. Front Cell Infect Microbiol 12:851784.

49. Smith P, Cozart J, Lynn BK, Alberts E, Frangipani E, Schuster M. 2019. Bacterial Cheaters Evade Punishment by Cyanide. iScience 19:101–109.

50. Bertani G. 1951. Studies on lysogenesis. I. The mode of phage liberation by lysogenic Escherichia coli. J Bacteriol 62:293–300.

51. Shanks RM, Caiazza NC, Hinsa SM, Toutain CM, O’Toole GA. 2006. *Saccharomyces cerevisiae*-based molecular tool kit for manipulation of genes from gram-negative bacteria. Appl Environ Microbiol 72:5027–36.

52. Neidhardt FC, Bloch PL, Smith DF. 1974. Culture medium for *enterobacteria*. Journal of Bacteriology 119:736–747.

